# Sustained nitrogen loss in a symbiotic association of Comammox *Nitrospira* and Anammox bacteria

**DOI:** 10.1101/2020.10.12.336248

**Authors:** Ekaterina Y. Gottshall, Sam J. Bryson, Kathryn I. Cogert, Matthieu Landreau, Christopher J. Sedlacek, David A. Stahl, Holger Daims, Mari Winkler

## Abstract

The discovery of complete aerobic and anaerobic ammonia-oxidizing bacteria (Comammox and Anammox) significantly altered our understanding of the global nitrogen cycle. A high affinity for ammonia (*K*_m(app),NH3_ ≈ 63nM) and oxygen place the first described isolate, Comammox *Nitrospira inopinata* in the same trophic category as organisms such as some ammonia-oxidizing archaea. However, *N. inopinata* has a relatively low affinity for nitrite (*K*_m,NO2_ ≈ 449.2μM) suggesting it would be less competitive for nitrite than other nitrite-consuming aerobes and anaerobes. We examined the ecological relevance of the disparate substrate affinities by coupling it with Anammox (*Nitrospira inopinata* and *Brocadia anammoxidans*, respectively). Synthetic communities were established in hydrogel granules in which Comammox grew in the aerobic outer layer to provide Anammox with nitrite in the inner anoxic core to form dinitrogen gas. This spatial organization was confirmed with FISH imaging, supporting a mutualistic or commensal relationship. Successful co-habitation of Comammox *N. inopinata* and Anammox in synthetic granules broadens our limited understanding of the interplay between these two species and offers potential biotechnological applications to study any type of bacterial pairings in a systematic and reproducible manner.

## INTRODUCTION

Anaerobic ammonia-oxidizing (Anammox) bacteria occupy a unique niche due to their ability to anaerobically oxidize ammonia along with nitrite to form dinitrogen gas ^1–5^. They are widely distributed in oxygen depleted environments across both marine and freshwater habitats where they often partner with aerobic ammonia-oxidizing archaea (AOA) to obtain the nitrite required for growth ^6–9^. This association is thought to be favored in oligotrophic environments such as the Oxygen Minimum Zones (OMZ) ^9–12^ by the high affinities of the marine AOA for ammonia and oxygen concentrations (ranging from an extremely low K_m(app), NH3_ ≈ 3 nM for the marine *Nitrosopumilus maritimus* to a K_m(app), NH3_ ≈ 5 µM for the terrestrial moderate thermophile *Nitrosotenius uzonensis*) ^13–18^. The unique physiology of Anammox has also found utility in nitrogen removal from high ammonia wastewater, where they generally partner with putatively r-strategist ammonium-oxidizing bacteria (AOB) having generally lower affinities for ammonia (K_m(app), NH4+_ ≈ 500 µM NH_4_^+^) but much higher growth rates than the AOA ^15,18,19^. In these systems the organisms generally grow together as biogranules, in which growth of oxygen-consuming AOB near the surface functions to sustain an inner Anammox sub-oxic core ^20,21^. The discovery of complete ammonia-oxidizing *Nitrospira* species (Comammox) ^22,23^ having ammonia (K_m(app), NH3_ ≈ 63 nM) ^18^ and oxygen affinities likely near that of AOA, suggests that they may also couple with Anammox in low ammonia environments and have possible functions in nitrogen removal in other treatment applications.

Comammox organisms produce nitrate as a product of ammonia oxidation with nitrite as an intermediate ^22,23^. Species are common in many natural and engineered systems, including forest and agricultural soils, freshwater and brackish sediments, and waste and drinking water treatment plants ^24–30^. As yet, the biotic and abiotic factors controlling their abundance and activity are not well defined. However, more recent analyses suggest that *Nitrospira* can become the dominant ammonia oxidizer in Water Resources Recovery Facilities (WRRFs) operated at low oxygen, and so offer more energy efficient nitrification in systems designed to operate at low dissolved oxygen (DO) levels ^30,31^. Two Comammox species (*N. nitrosa* and *N. nitrificans*) were detected together with Anammox in an enrichment culture. The co-occurrence of these Comammox organisms and *Brocadia* suggested a functional relationship between these two organisms ^23^. Comammox was also found to coexist with AOB and Anammox in a sequencing batch reactor for sludge digester liquor treatment and responsible for ∼25% nitrogen removal ^32^.

Mathematical models have indicated that oxygen and ammonium concentrations are major niche differentiating factors for Anammox and AOA or Anammox and AOB. For example, the association between AOB/AOA and Anammox is often disrupted by nitrite-oxidizing bacteria (NOB) competing with Anammox for nitrite ^33,34^, resulting in a lower conversion of ammonia to N_2_ and an increase in nitrate formation. Thus, there is clear need to resolve competitive and cooperative relationships among Comammox, Anammox, AOA, AOB, and NOB in order to develop a better predictive understanding of their roles in the global nitrogen cycle and to better define possible applications in engineered systems ^12^. The Comammox relationship with Anammox is of specific interest since both bacteria utilize ammonia and nitrite and are active in low DO environments. However, comparative laboratory analyses of organisms that function cooperatively at the boundary between oxic and anoxic environments have been limited for lack of appropriate experimental systems. Suspended microbial cultures do not form the oxygen gradient essential for partnership. Chemostat studies must balance the conflicting nutrient requirements for cooperative growth and are prone to washout. We here use synthetic community assemblies of Comammox *N. inopinata* and Anammox (dominant species *Brocadia anammoxidans*) immobilized in hydrogel granules to evaluate environmental conditions supporting their partnership.

The hydrogel format, which is a three-dimensional network of hydrophilic polymers, has been successfully implemented to immobilize active nitrifying and denitrifying bacteria ^35^, denitrifying cultures ^36^, anaerobes originating from hydrothermal vents ^37^, and Anammox ^38^. The gels can also be formed to resemble naturally occurring biogranules ^39,40^, in which Comammox and Anammox can be entrapped in a matrix of extracellular polymeric substances (EPS) that promotes the formation of stable nutrient gradients and microbial interactions on a micrometer scale. Using a synthetic hydrogel set-up to achieve conditions for both aerobic and anaerobic growth, we now demonstrate the formation of a stable partnership in which Comammox competitively excludes the AOB to supply nitrite to Anammox and, in turn, Anammox lowers nitrite to a non-inhibitory concentration. This is the first clear evidence for possible ecological relevance of an association that should be useful for various biotechnological applications.

## RESULTS

### Comammox and Anammox bacteria establish cooperation within synthetic granules in batch cultures

A hydrogel formulation was developed to entrap high numbers of the only available Comammox pure culture *N. inopinata* and an Anammox enrichment (dominant organism *Brocadia anammoxidans*) in synthetic Polyvinyl alcohol – sodium alginate (PVA-SA) hydrogel beads. Comammox-Anammox beads demonstrated a constant rate of ammonium consumption over the study period of 86 days, resulting in complete nitrogen removal from the system (Fig. 1a). Since no external nitrite was supplied, Comammox activity was the sole source of nitrite for Anammox. We did not observe measurable nitrite or nitrate accumulation throughout the incubation period as would be expected from Comammox ^22,23^ or Anammox ^2,41^ metabolism alone. Beads containing only Comammox stoichiometrically oxidized ammonia to nitrate, with transient accumulation of nitrite (Fig. 1b), and beads containing only the Anammox enrichment converted ammonium to N_2_ when supplied with ammonia and nitrite at the ratio of 1:1.3 (Fig. 1c). Comammox-Anammox bead consumption of 1mM ammonium was about 3-fold slower than observed for beads containing only Comammox (0.021 mM NH_4_/day versus 0.064 mM NH_4_/day).

**Figure 1.**
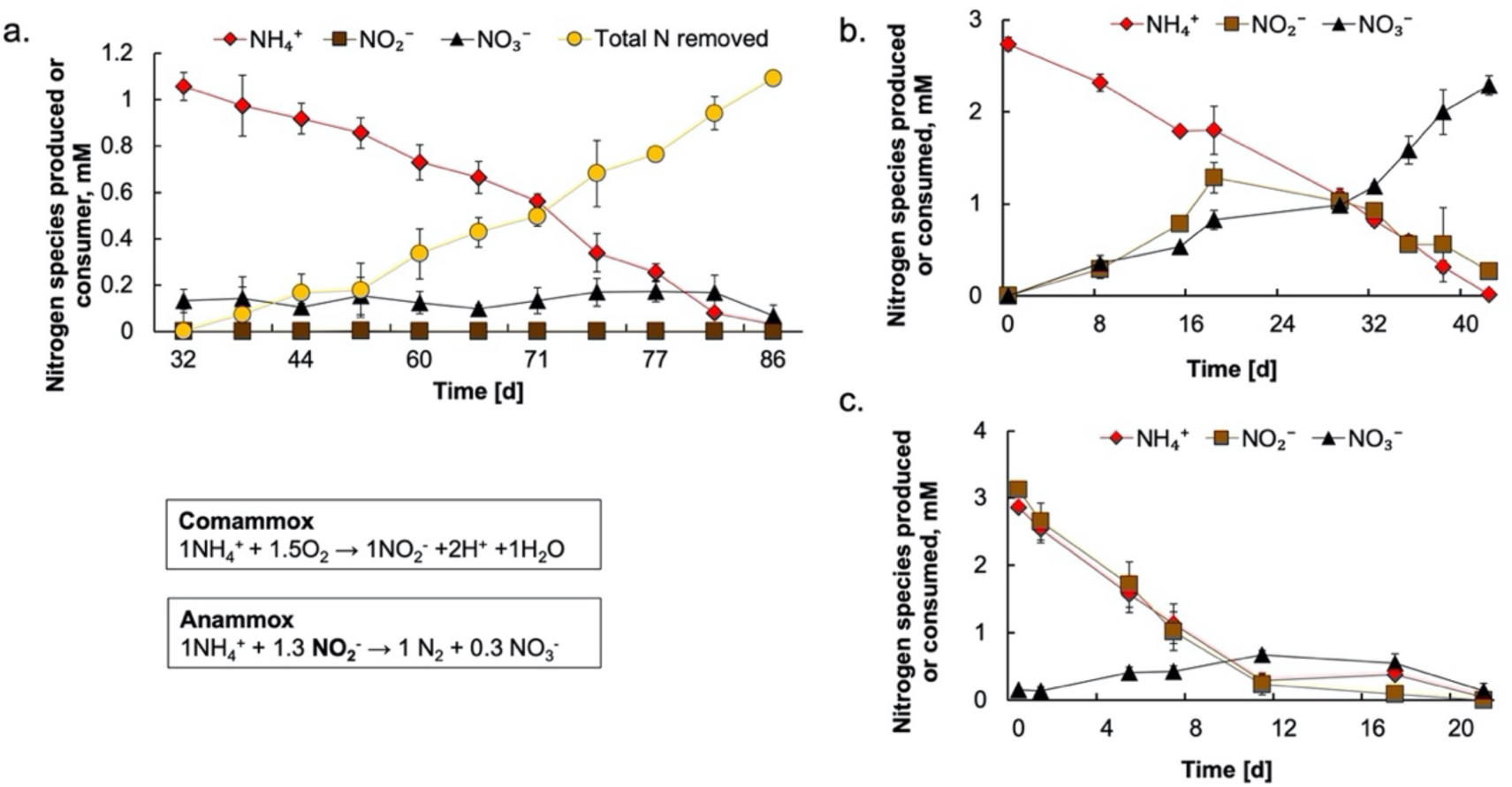
Comammox and Anammox cooperation within hydrogel beads. (a) Immobilized Comammox and Anammox hydrogel beads demonstrated cooperation at 4 mgO_2_/L and with ammonium as sole source of energy, electrons, and nitrogen in three biological replicates (n=3). (b) In the absence of Anammox the gel immobilized Comammox was capable of further oxidizing nitrite to nitrate. (c) In the absence of Comammox, the established Anammox anabolic reaction pathway was demonstrated.

### qPCR supports Anammox and Comammox cooperation in hydrogel beads

Total bacterial abundance, as measured by qPCR targeting the 16S rRNA gene, ranged from 10^5^ to 10^6^ gene copies ng^-1^ of granule material sampled at the beginning and the end of the incubation period from both entrapped and planktonic communities (Suppl. Fig. 3). Quantification of the *amoA* of Comammox and betaproteobacterial AOB, and of Anammox 16S rRNA genes, demonstrated that Comammox and Anammox bacteria dominated the ammonia-oxidizer population in the hydrogel beads (Fig. 2). In contrast, betaproteobacterial AOB were near the limit of detection in all hydrogel samples and persisted only in the fraction of non-immobilized planktonic bacteria (Fig. 2).

**Figure 2.**
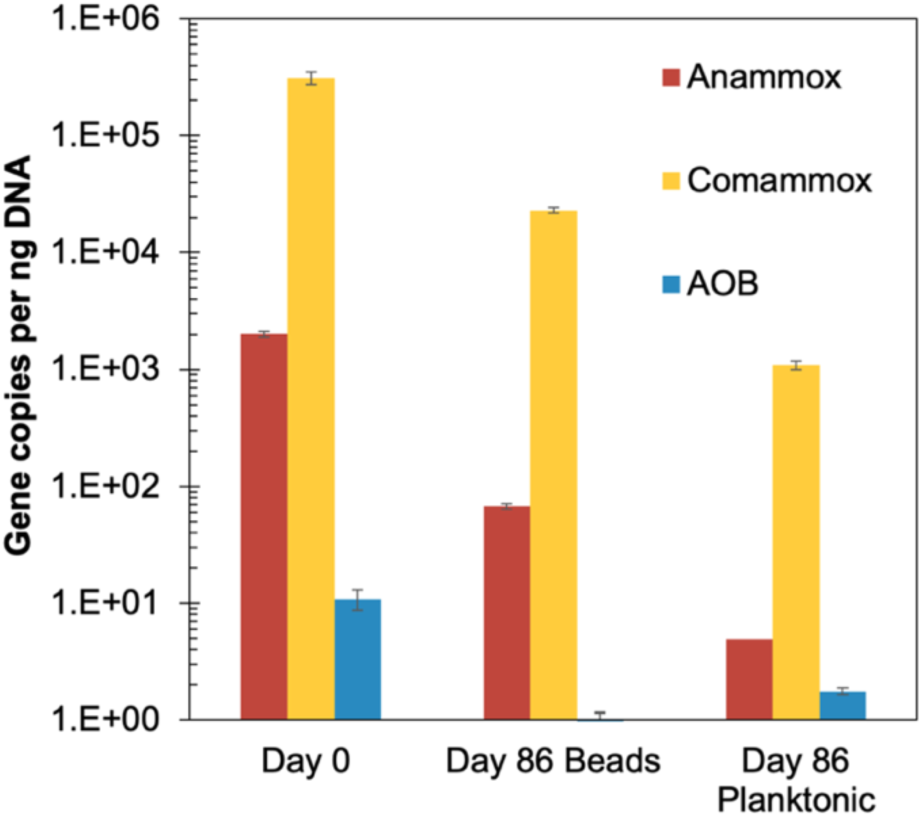
Abundance of Comammox, Anammox, and canonical AOB in hydrogel beads and in the planktonic fraction determined by qPCR of *amoA* genes (Comammox, AOB) and 16S rRNA genes (Anammox). Gene copy numbers per ng of total extracted DNA are shown for samples taken at the start of the incubation (day 0) and after 86 days. Genetic material corresponding to the initial biomass (day 0) was obtained directly after immobilization through the same extraction method as the day 86 biomass samples. Gene copy numbers were adjusted according to their average occurrence (Supplementary Table 3).

**Figure 3.**
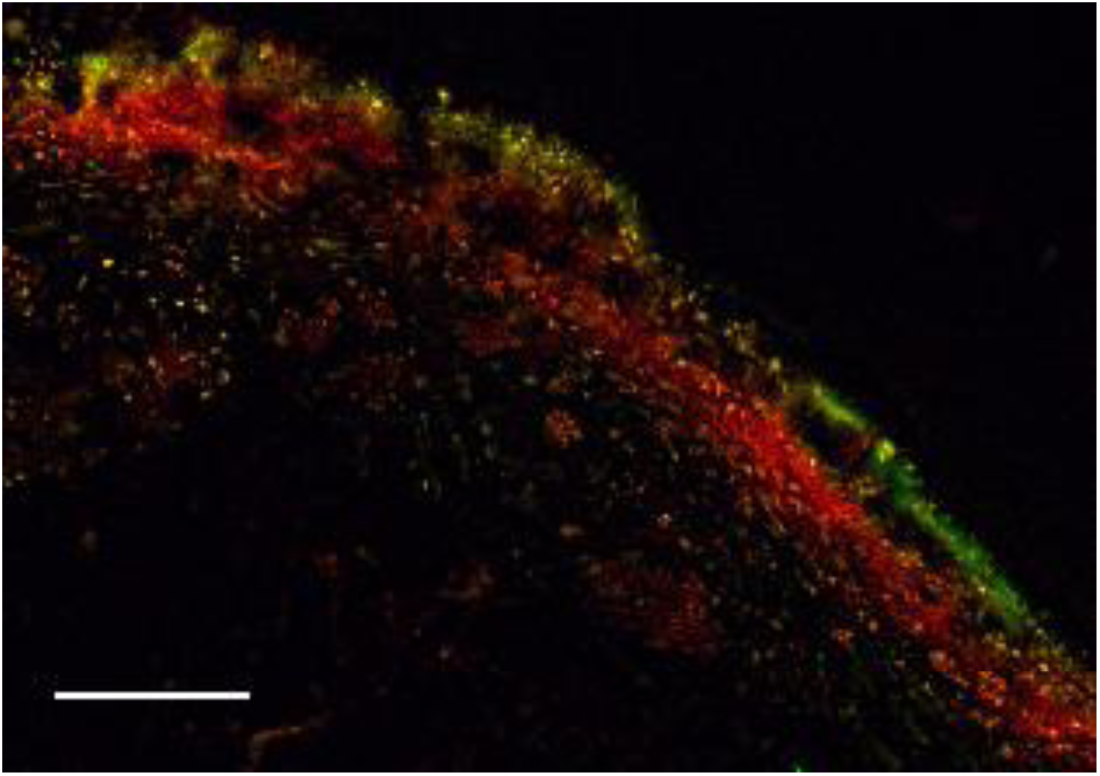
Comammox *Nitrospira* and Anammox demonstrate spatial segregation within hydrogel beads. Simultaneous *in situ* hybridization of a bead section with Cy3-labeled probe AMX368 (red; Anammox), fluorescein-labeled probe Ntsp662 (green; *Nitrospira*), and Cy5-labeled probe Nso1225 (blue; betaproteobacterial AOB). The bar represents 100 μm.

### 16S rRNA sequencing demonstrated Comammox N. inopinata and Anammox presence after the incubation period while AOB and NOB remained low but constant

The community composition of PVA-SA hydrogels at the beginning and end of incubation was investigated using amplicon sequencing of the V4-V5 region of the 16S rRNA gene (Suppl. Fig. 1). In total 409 operational taxonomic units (OTUs) were identified. Relative abundance of reads assigned to OTUs related to heterotrophic organisms increased by 22.6% from beginning to end of incubation (Suppl. Fig. 1). There were corresponding shifts in relative abundances of OTUs assigned to ‘*Candidatus Brocadiaceae’* (Anammox), *Nitrospira inopinata* (Comammox), *Nitrosomonas* (AOB), *Nitrosopumilis* or *Nitrososphaera* (AOA), and other *Nitrospira* (NOB) that were not assigned to the *Nitrospira inopinata* OTU between the initial biomass and after 86 days (Suppl. Fig. 2). The Comammox OTU accounted for 20.3% reads at the beginning of the incubation and averaged 5.15% (± 7.73% S.D., n=3) at termination. Canonical AOB, AOA, and NOB accounted for only 0.060%, 0.010%, and 0.020% of the reads initially and remained low throughout the experimental period, with final average relative abundances of 0.010%, 0.017%, and 0.0033% respectively (S.D. ≤ 0.01%) after 86 days of cultivation under nitrifying conditions (Suppl. Fig 2). A single OTU assigned to *N. inopinata* was the only nitrifier OTU of substantial relative abundance. Thus, it can be inferred that it alone was responsible for the observed nitrification activity (Fig.1a-b). Together this indicated that Comammox was responsible for partial nitrification in support of Anammox. The observed apparent loss of Anammox in PVA-SA hydrogels after 86 days, as represented by 7.68% of the reads in the initial beads and 0.32% (± 0.05% S.D., n=3) of the final biomass (Suppl. Fig. 2), likely reflected a combination of degradative loss in the nutrient-limited interior and by oxygen exposure near the outer boundary of the granules, increased heterotroph abundance, and poor extraction of DNA from Anammox cells embedded deep within hydrogel granules. Additionally, PCR biases for the v4-v5 16S rRNA gene primers may preferentially amplify other taxa that increased in abundance during incubation, skewing relative abundance data ^42^.

### Comammox and Anammox species are spatially segregated within hydrogel beads

Given the efficient removal of nitrogen by this community, we further examined the population structure using rRNA-targeted FISH imaging and qPCR. Fluorescent probe AMX368-hybridized Anammox bacteria were detected in the initial bead samples as well as at the end of the incubation period (Fig. 3), confirming Anammox persistence within hydrogel beads. Following initial bead fabrication, cells were dispersed in the polymer matrix while they mainly appeared on the peripheral layer of the beads after 86 days of incubation (Fig. 3). While the Comammox organisms *N. inopinata* (green) was randomly distributed just after immobilization, it formed a thin layer at the surface following the incubation period. In particular, Comammox cells resided at the oxygenated edge of the hydrogel beads, whereas Anammox cells were situated in a deeper (presumably anoxic) layer. Probe Nso1225 (specific for betaproteobacterial AOB) did not produce a detectable fluorescent signal (Fig. 3).

## DISCUSSION

The ability of Comammox microorganisms to survive under low substrate concentrations ^18^ allows them to occupy oligotrophic environments. Cooperation with Anammox bacteria has been reported in bioreactor studies ^23,43^. Here we used a unique hydrogel format to pair Comammox and Anammox in a granule-like structure engineered to promote the formation of nutrient gradients theorized to foster their co-occurrence within natural habitats. Microbial activities in natural biofilms or biogranules generate substrate gradients, allowing aerobic organisms (like Comammox) to grow on the biofilm periphery and anaerobes (like Anammox) in the oxygen depleted adjacent inner layers ^44,45^. The hydrogel format promotes a similar relationship in a controlled laboratory setting. Therefore, we fabricated synthetic hydrogel granules harboring a single species of Comammox, *Nitrospira inopinata*, and an enrichment of Anammox related to *Brocardia*. The beads remained intact throughout the 86-day incubation period, as previously shown in our study of demonstrating retention of active AOA in hydrogel constructs ^46^. Cooperation was confirmed by a nearly total nitrogen removal and negligible production of nitrate (Fig. 1).

The high total nitrogen removal was consistent with Anammox being sustained by nitrite produced by the co-immobilized *N. inopinata*, suggesting that very little nitrite was further oxidized to nitrate in the synthetic community. Low nitrate accumulation sets the cooperation described here apart from previously studied systems of AOB/AOA and Anammox or AOB/AOA and NOB, in which significant nitrate was produced. A Comammox-Anammox partnership may therefore be beneficial in engineering applications that aim to eliminate nitrogen species using autotrophic microbial populations. While the NH_3_ affinity of AOB is moderate to low, AOA and Comammox have high affinities for NH_3_, which is beneficial for better N removal in mainstream WRRFs. However, AOA are generally less abundant in WRRFs possibly due to organic materials inhibition leading to poor bioavailability of copper in these settings ^47^. Comammox, in contrast, can be quite prevalent in WRRFs ^28,30^. From this perspective, Comammox may be better adapted than AOA for engineered systems and could therefore be paired with Anammox for N removal in those systems.

Together these results suggest that nitrite was consumed by Anammox and not further oxidized by Comammox. This is in alignment with reported nitrite affinities of *K*_m(app), NO2_ ≈ 48 µM for Anammox and *K*_m(app), NO2_ ≈ 449.2 μM for *N. inopinata*, suggesting that a higher rate of nitrite consumption by Anammox provided it a competitive advantage. We also determined a nitrite half-inhibition constant of *Ki*_,NO2_=0.31 ± 0.19 mM nitrite on *N. inopinata* growth (µMax= 2.6 ± 0.75) and no inhibition at low concentrations (<0.1 mM) (Suppl. Fig. 4). This suggests that Anammox serves Comammox by keeping nitrite concentrations at very low and non-inhibitory levels. In turn, Comammox serves Anammox by consuming oxygen and providing an anoxic environment while also supplying nitrite. Thus, maintaining a non-inhibitory concentration of nitrite in the Comammox local environment by Anammox appears to foster a symbiotic relationship between the two microorganisms. However, a novel Comammox species, *Nitrospira kreftii*, was recently identified displaying physiological characteristics distinct from *N. inopinata*. Particularly, *N. kreftii* has a higher nitrite affinity (*K*_m(app), NO2_ ≈ 13 µM) altering the niche determination understanding of Comammox and its potential interactions with other nitrifiers ^48^. Thus, for specific engineering solutions, it is important to understand the kinetic parameters of different Comammox species.

In order to confirm that observed ammonia oxidation and nitrogen removal resulted from the activity of Comammox and Anammox organisms, and not from other ammonia oxidizers, we conducted a population analysis by sequencing the hydrogel beads samples. 16S rRNA analysis of microbial populations from the initial beads versus those incubated for 86 days beads revealed that *N. inopinata* remained the major aerobic ammonia oxidizer. Other nitrifying organisms AOA, AOB, and NOB accounted for only a small fraction of the OTUs (<0.06%) and slightly decreased in relative abundance after the incubation of 86 days. (Fig. 3).

This data contrast with a previous study of a sequencing batch reactor reporting relative abundances of AOB (18%), Anammox (0.1%), and Comammox (0.2%) during reactor operation^32^. The high abundance of canonical AOB (18% vs. less than 1 % of Comammox) did not fully support the proposed cooperation between Comammox and Anammox. In our study, however, Comammox was demonstrated to be responsible for partial nitrification in the constructed hydrogel beads, as deduced from the virtual absence of other known ammonia-oxidizing organisms. However, the presence of heterotrophic populations originating from the Anammox enrichment culture (Suppl. Fig. 1) may also have been of functional significance.

Members of the *Ignavibacteria* taxa, some of which are capable of performing dissimilatory nitrate reduction to ammonia (DNRA) under anoxic conditions ^49^, remained stable throughout the batch test. Also of possible relevance to nitrogen loss was the activity of populations assigned to four proteobacterial orders, *Xanthomonadales, Burkholderiales, Rhodocyclales*, and *Rhizobiales*, all of which are known to contain species capable of facultative anaerobic heterotrophic denitrification ^50^. Low nitrate concentrations in the effluent might in part be attributed to the activity of heterotrophic denitrifiers, relying on organic carbon from degradation of alginate, biomass decay ^51^, or small organic acids produced by autotrophic organisms ^52,53^. Alginate did not appear to provide a significant source of carbon since bead integrity remained stable throughout the study period and the observed nitrate consumption was inconsistent with alginate being a significant carbon source for denitrifying heterotrophs (Suppl. Table. 1). Although it is unclear what nutrients supported heterotopic growth, the high abundance of aerobic heterotrophs suggest that oxygen and not nitrate served as the primary terminal electron acceptor, which would lead to a competition between heterotrophs and *N. inopinata* for oxygen in the outer edges of the hydrogel granules, and utilizing nitrate (if any) within the anaerobic core volume of the granules.

In addition to confirming by qPCR the retention of Comammox and Anammox, and an insignificant contribution of AOB and NOB (Fig. 2), we demonstrated their expected spatial relationship using fluorescent probes specific to the associated bacterial families (Fig. 3). The fluorescent Ntsp662 oligonucleotide probe specific to *Nitrospira* showed that Comammox cells were most abundant in the outer layer of the beads, as dictated by the oxygen dependence of aerobic ammonia oxidation. The fewer cells observed deeper in the granule interior may be an artifact of introducing bacteria into interior during the slicing procedure or possibly reflecting the ability of Comammox to grow at lower substrate concentrations typically present in deeper regions and inactive Comammox retaining high ribosome content detectable by FISH. The Anammox bacteria were detected using fluorescent Amx368 probe and were distributed in small aggregates throughout the inner core space but mostly clustered immediately below Comammox towards the edge of the bead where substrate supply is highest. Predicted high affinity for oxygen therefore can explain Comammox spatial localization within the outer layer of the hydrogel beads and supports a cooperative relationship of Comammox and Anammox in sub-oxic environments.

In conclusion, we demonstrated that the recently described complete ammonia-oxidizing organism *N. inopinata* can establish a cooperative relationship with anaerobic ammonia-oxidizing bacteria in a synthetic granule format. The significance of this novel cooperation was documented by the low nitrate formation and the competitiveness of the Comammox organism over other aerobic ammonia- and nitrite-oxidizers. The stability of this association is suggestive of other interactions, such as the release of small organic compounds (e.g., formate) from autotrophic Anammox ^54–56^ serving as electron donors for heterotrophs or possible mixotrophic lifestyle of Comammox. Since nitrate is an undesirable product of some engineered treatment systems, the Comammox-Anammox symbiosis may be of economic and ecological importance to reduce nitrogen contamination of receiving waters. In addition, our study also suggests that a similar cooperation of Comammox and Anammox may be important in natural aquatic or terrestrial habitats that are oxygen and ammonia deplete.

## METHODS

### Microbial strains and growth

Comammox *Nitrospira inopinata* strain was incubated in limited mineral media aerobically at 37°C in the presence of 1mM NH_4_^+ 22^. Anammox biomass was obtained from a wastewater treatment plant in Rotterdam Sluisjesdijk (the Netherlands) and was maintained anaerobically in a plug-flow glass column supplied with 1mM NH_4_^+^ and 1.3mM NO_2_^-^ and operated at 30°C. The dominant Anammox organism in this sludge was reported to be *Brocadia anammoxidans* ^57^. The mineral media used for Anammox growth had following composition ^41,58^: 2.5 mM MgSO_4_·7H_2_O, 1.2 mM CaCl_2_, 5 mM NaHCO_3_, 0.2 mM KH_2_PO_4_, FeSO_4_·7H_2_O 9.0 mg, EDTA·2Na 5 mg. Trace elements solution (per 1L ddH_2_O): EDTA, 15 g; ZnSO_4_·7H_2_O, 0.43 g; CoCl_2_·6H_2_O, 0.24 g; MnCl_2_. 4H_2_O, 0.99 g; CuSO_4_·5H_2_O, 0.25 g; NaMoO_4_·2H_2_O, 0.22 g; NiCl_2_·6H2O, 0.19 g; NaSeO_4_·10H_2_O, 0.21 g; H_3_BO_4_, 0.014 g.

### Hydrogel beads fabrication

A 10% Polyvinyl alcohol (PVA) (w/v) and 2% Sodium alginate (SA) (w/v) solution in water was prepared and sterilized as previously described ^46^. Comammox *Nitrospira inopinata* culture was concentrated 100-fold via 0.1 µm tangential filtration cassette (Millipore Sigma). Comammox viability after filtration was assessed via incubation in the mineral medium with subsequent quantification of nitrification activity. Granules enriched in Anammox were disaggregated using a blender for 2 minutes and resuspended in the mineral medium. Comammox and Anammox bacteria were mixed in a 1:1 ratio and then interspersed with the PVA-SA polymer solution to achieve 6% PVA and 1% SA concentrations. Using the 0.1 mm diameter tips PVA-SA-bacteria mixture was dropped into a 2% (w/v) Barium chloride (BaCl_2_) bath in order to form spherical hydrogel particles. The hydrogel beads were allowed to harden in the BaCl_2_ solution for 1 hour before rinsing several times in a sodium chloride solution (0.9%) to remove excess polymer. All solutions and medium used for the hydrogel bead production were prepared anaerobically to minimize Anammox exposure to oxygen. The granules were incubated in batch bottles containing an ammonia-mineral media ^22^ without organic carbon and their activity was monitored for 86 days. Three sets of Comammox-Anammox beads were set up as biological replicas and incubated in parallel. The co-culture was oxygenated with ca. 4 mgO_2_/L 3 times per week and ammonium was replenished when depleted. The microbial activity was measured using spectrophotometric analysis of nitrogen species concentration in the growth media throughout the incubation period. Two additional batch bottles were set up to separately assess the independent growth of Comammox and Anammox bacteria, respectively (Suppl. Fig 1 and 2). Dissolved oxygen concentration was monitored with oxygen sensor spots (PyroScience GmbH, OXSP5).

### Microbial activity measurements

Ammonium (NH_4_^+^), nitrite (NO_2_^-^), and nitrate (NO_3_^-^) concentrations were measured using a photometric method with Gallery^™^ Automated Photometric Analyzer (ThermoFisher Scientific, Waltham, MA U.S.A.) with the Total Oxidized Nitrogen (TON)-Nitrite and Ammonia reagents calibrated using Sodium nitrite (NaNO_2_) and ammonium chloride (NH_4_Cl) standards. Samples were processed immediately after collection.

### Molecular analysis of microbial community composition

Genomic DNA was extracted from PVA-SA hydrogel beads by bead-beating and from growth media (two series of 25 seconds each, Bullet Blender 5 homogenizer) and subsequent purification using a Power Biofilm kit (DNeasy Power Biofilm kit, Qiagen) following manufacturer instructions. Purified DNA samples were diluted to 20 ng/µl and sent to MR DNA (Shallowater, USA) for construction of amplicon libraries for the V4-V5 region of 16S rRNA genes using primers 515F-Y (5′-GTGYCAGCMGCCGCGGTAA) ^59^ and 926R (5’-CCGYCAATTYMTTTRAGTTT) ^60^. Sequencing was performed on an Illumina MiSeq producing 300 bp paired-end reads. Sequence libraries were processed and an OTU table was constructed using the USEARCH v11 package ^61^. Forward and reverse reads were merged and demultiplexed with default settings, yielding 15,837, 13,628, 21,460, and 20,563 reads for the samples from the initial biomass and the three batch replicates, respectively. OTU selection was performed by quality filtering these reads with “-fastq_maxee=1.0”, dereplicating with “minuniquesize=20” and running “unoise3” in “zotu mode with default settings ^62^. Taxonomy was assigned using the sintax command ^63^ and the RDP 16S training database (release 11) ^64^. OTU counts were determined by mapping all merged reads then rarefied to 10,000 reads per sample.

### Fluorescence in situ hybridization

Fixation and hybridization of the bacterial hydrogel beads samples was carried as previously described ^65^ with the following modifications. Hydrogel beads were fixed in 4% paraformaldehyde for 3-16 hours at 4°C and washed with Ethanol/PBS (50:50 w/v) three times. If not processed immediately, samples were stored in PBS/Ethanol 50:50 solution at −20°C. Fixed hydrogel beads were submerged in OCT Compound (Agar Scientific) or NEG-50 ^™^ (Richard-Allan Scientific ^™^) overnight and then frozen at −20°C for 3-16 hours prior to sectioning. Thin 20-30 nm sections of hydrogel beads were produced using a micro-cryotome (Cryostar NX50™) at −20°C and mounted on gelatin coated Teflon-covered glass slides. Thin sections were dried in the oven at 46°C for 10 minutes, sequentially dehydrated in 50, 80, 95% ethanol, air dried, and stored at −20°C. Using Liquid Blocker Pap Pen (Life Technologies, Carlsbad, CA, USA) a hydrophobic barrier was applied to glass slides containing thin cryosections. Oligonucleotide probes are summarized in Suppl. Table 4 and were applied in equimolar concentration to the thin sections and incubated at 46°C for 2 hours. All probes were purchased from Eurofins, Louisville, KY. The slides were mounted in the Vectashield antifade mounting medium (Cole-Parmer). Confocal laser scanning microscope Zeiss LSM700 (Carl Zeiss, Germany) equipped with an Axio cam 503 mono camera was used to visualize hybridized cryosections with 488, 543, and 633 nm lasers. A 10x/0.45 objective was used to image large regions of the sections. 40x/1.3 and 100×/1.46 Oil Ph3 plan-apochromat oil objectives were used in order to obtain higher-resolution images. The images were collected in the Zen Blue software.

### Quantitative PCR

In order to quantify Comammox, Anammox, AOB, and NOB in hydrogel beads a quantitative PCR (qPCR) analysis was performed. Specific primers are listed in Supplementary Table 2. All qPCR assays used thermocycling conditions described in the reference articles (Suppl. Table 2) and were performed on Roche LightCycler 480 high-throughput real-time PCR system in white LightCycler 480 multiwell tubes and associated LightCycler 480 transparent caps (Roche Molecular Systems, Inc, Pleasanton, CA, United States). Triplicate reactions were performed for each sample. Amplification was performed according to protocols described by Fowler et al. (2018) ^24^. Briefly, each reaction contained 5 μl Master Mix (DreamTaq Green PCR Master Mix, Thermo Scientific), 3 μl nuclease free water, 1 μl sample DNA (final reaction contained 1-2.5 ng of DNA), 0.5 μl of each PCR primers (10 μM) added to a final volume of 10 μL per reaction. Denaturation was performed at 95°C for 5 min followed by 35 cycles of 15 s denaturation at 95°C, 15 sec annealing at target gene temperature, and 13 sec extension at 72°C. After each qPCR assay, the specificity of the amplification was tested with melting curve analysis and agarose gel electrophoresis.

The gene copy number for the standards was calculated from DNA concentration obtained using Qubit double stranded DNA High Sensitivity Assay kit (Thermo Fisher Scientific). Ten-fold series dilutions were prepared for each standard over five orders of magnitude from 10^1^ to 10^5^ copies and were amplified in triplicates. For use as a qPCR standard for Anammox, a plasmid was constructed by cloning the 808 to 1066 16S rRNA gene region using 808F ^66^ and 1066R ^67^into the pCR™4-TOPO® TA vector (TOPO® TA Cloning® Kit, InvitrogenTM) and sequencing (Eurofins Scientific) was performed for validation. qPCR was conducted using the 818F ^67^and 1040R ^66^ primers. Gene copy numbers per ng of DNA were adjusted according to the reported gene copy numbers (Suppl. Table 3).

### Nitrite inhibition assay

Strains were grown in a mineral salts medium modified from Bollmann et al. (2011) to include the trace elements and Ethylenediaminetetraacetic acid ferric sodium salt typically used in saltwater AOA medium ^68^. Nitrogen species were measured daily using the Thermo Fisher Gallery Analyzer™. Nitrate production rate was measured under varying nitrite concentrations (0.1mM, 0.25, mM, 0.5 mM, 0.6mM, 0.75 mM, 2.5 mM, 9mM) in the presence of 1 mM ammonium in individual 100-mL batch cultures. Daily ammonium, nitrite, and nitrate measurements were made until either 90% of ammonium had been consumed and the culture was considered to be in late exponential phase or two weeks had passed, whichever occurred first. Kinetic constants were fit to a typical inhibition curve with the non-linear least squares model in the R stats package.

## ACKNOWLEDGEMENTS

We thank Bao N. Quoc and Dr. Kristopher A. Hunt for helpful discussion and theoretical calculation assistance. EYG, SJB, KIC, ML, DAS, and MW received financial support from the Defense Advanced Research Projects Agency [Contract Number: HR0011-17-2-0064] and U.S. Department of Energy (DOE) [Contract Number: DE-SC0020356]. CJS and HD were supported by the Austrian Science Fund (project P30570-B29). SJB received support from the Washington Research Foundation.

## AUTHORS CONTRIBUTIONS

EYG, SJB, KIC, ML, DAS, and MW designed the experiments. EYG, SJB, KIC, and ML performed the experiments and analyzed the data. EYG, SJB, KIC, ML, DAS, MW, CJS, and HD wrote the manuscript. Al authors read and approved the final manuscript.

## DATA AVAILABILITY

The authors state that all data supporting the findings of this study is available within the manuscript, and the supplementary materials files, or from the corresponding author per request.

