## Supplementary Table 2, Suppl. Fig. 2 for "Sustained nitrogen loss in a symbiotic association of Comammox *Nitrospira* and Anammox bacteria"

Gottshall E. *et al*.

**This file contains:**

Supplementary Tables 1 to 4
Supplementary Figs. 1 to 4

**Supplementary Table 1. Theoretically bioavailable carbon and nitrate in hydrogel beads**

| |
|---|
| Sodium alginate theoretically available per vessel, electron mol: |
| $$0.3\ g\ \cdot \frac{mol}{216.12\ g} \cdot \frac{20\ electrons}{mol} = \frac{0.02776\ electrons}{vessel}$$ |
| Nitrate theoretically available per vessel, electron mol: |
| $$30\ mg\ NO3\ \cdot \frac{1\ mmol\ NO3}{14\ mg\ \cdot 1L}\ \cdot\ 0.07L \cdot \frac{5\ electrons}{1\ mol\ NO3} \cdot \frac{1\ mol\ NO3}{1000\ mmol} = \frac{0.00075\ electrons}{vessel}$$ |

**Supplementary Table 2. 16S rRNA oligonucleotide probes**

| Target gene | Primer name | Forward primer 5'-3' | Reverse primer 3'-5' | Reference |
|---|---|---|---|---|
| Ca. *Nitrospira inopinata amoA* | Inopinata amoA-410F/815R | TCACCTTGTTGCTAACTAGAAACTGG | TCCGCGTGAGCCAATGT | [1] |
| Anammox 16S rRNA | 818F/ 1040R | ATGGGCACTMRGTAGAGGGGTTT | CAGCCATGCAACACCTGTRATA | [2,3] |
| AOB *amoA* | amoA-1F/2R | GGGG TTTCTACTGGTGGT | CCCCTCKGSAAAGCCTTCTTC | [4] |
| NOB *nxrB* | 169f /638R | TAC ATG TGG TGG AAC A | CGG TTC TGG TCR ATC A | [5] |
| Total bacteria 16S rRNA | 515F-Y/926R | GTGYCAGCMGCCGCGGTAA | CCGYCAATTYMTTTRAGTTT | [6] |

**Supplementary Table 3. Gene copy numbers for target genes**

| **Gene** | **Copy number** | **Reference** |
|---|---|---|
| Comammox *amoA* | 1 | [7] |
| AOB *amoA* | 3 | [8] |
| Anammox 16S rRNA | 1 | [9] |
| Heterotrophic 16S rRNA | ~ 3.8 | [10] |

**Supplementary Table 4. FISH probes**

| Probe name | Target organisms | Formamide, % | Reference |
|---|---|---|---|
| Ntspa662 | *Nitrospira* | 35 | [11] |
| Amx368 | All Anammox bacteria | 35 | [12] |
| Nso1225 | Betaproteobacterial ammonia-oxidizing bacteria | 35 | [13] |
| Nso192 | *Nitrosomonas oligotropha* lineage Cluster 6a | 35 | [14] |
| Eub338 | Most bacteria | 35 | [15] |

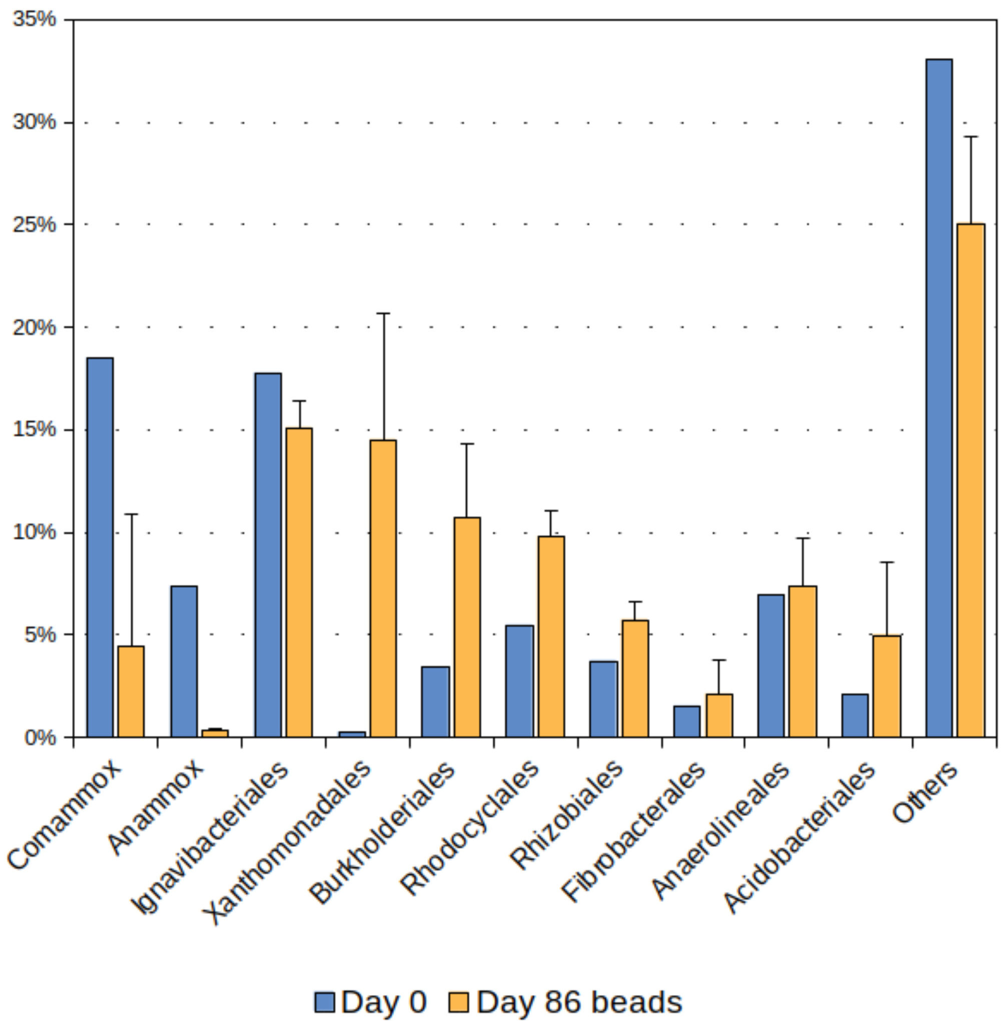


**Supplementary Figure 1**. Relative abundance of main microbial taxa within Comammox-Anammox beads as determined by sequencing of the V4-V5 region of the 16S rRNA gene. The columns from left to right correspond to the combined relative abundances of reads mapped to single OTUs with > 1% in any of the samples (Anammox was represented by five OTUs). “Others” refers to unclassified OTUs and taxonomic orders that did not have representative OTUs with greater than 1% relative abundance. “Day 86 beads” depicts the average of three replicates (n=3) with standard deviation presented as the positive error bars.

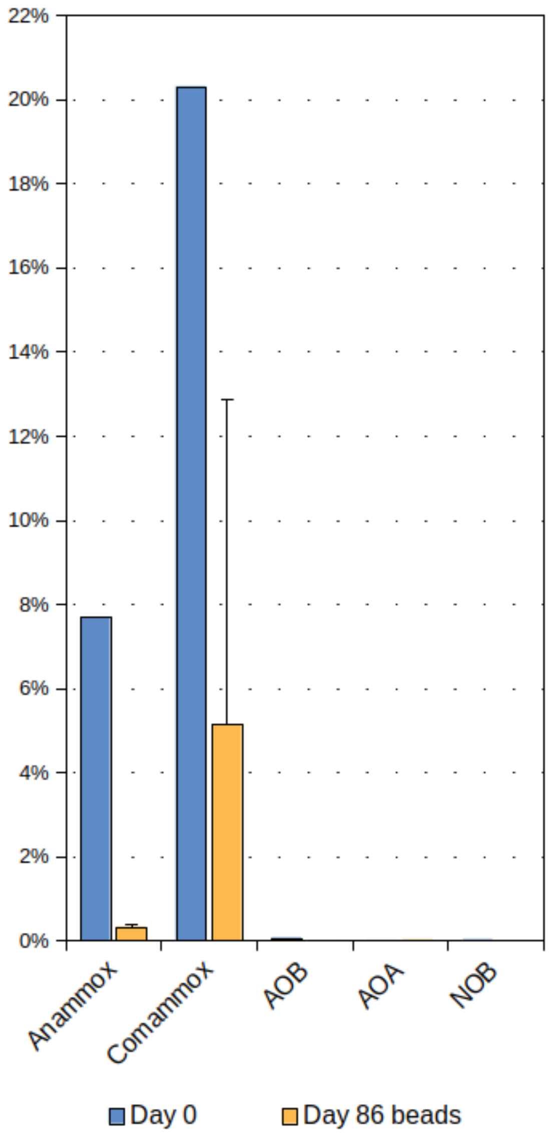


**Supplementary Figure 2.** Percent relative abundance for reads assigned to OTUs for '*Candidatus Brocadiaceae'* (Anammox), *Nitrospira inopinata* (Comammox), ammonia-oxidizing bacteria (AOB), ammonia-oxidizing archaea (AOA), and nitrite-oxidizing bacteria (NOB). Relative abundances for the three samples of beads (n=3) taken at Day 86 are averaged and the positive error bar indicates the standard deviation of the replicates. Comammox *Nitrospira* and Anammox *Brocadia* remained significantly present in the hydrogel beads while AOB, AOA, and NOB species remained consistently low.

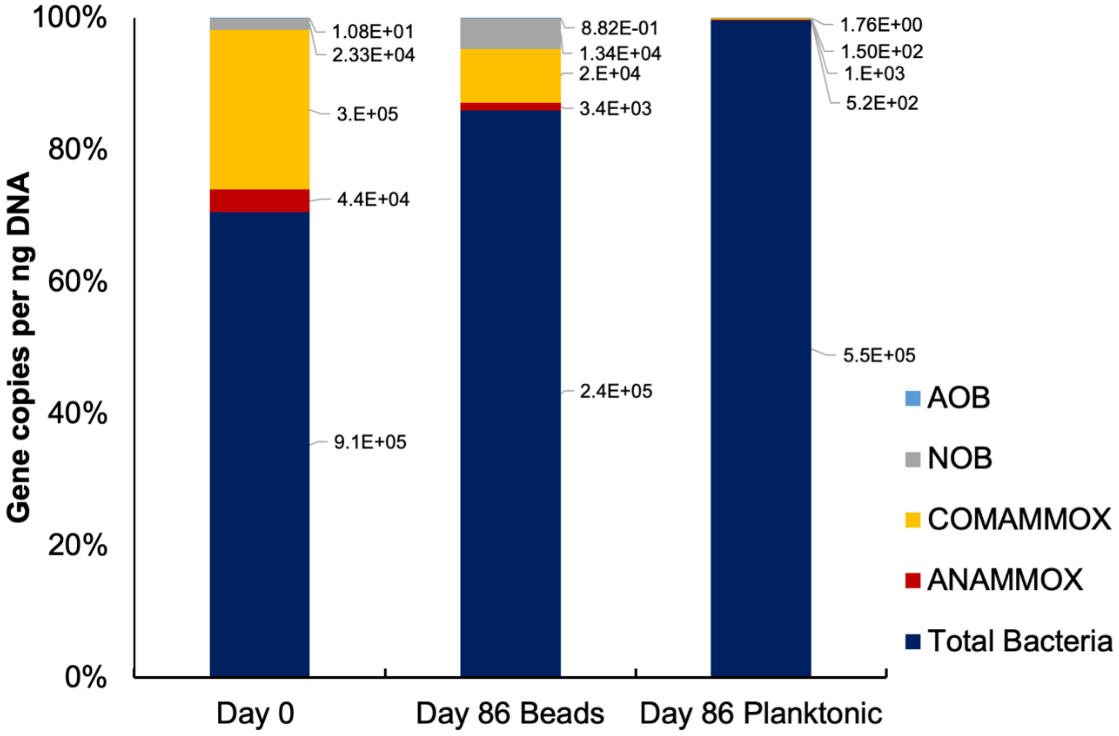


**Supplementary Figure 3**. Abundance of Total Bacteria, Comammox, Anammox, NOB, and canonical AOB in hydrogel beads and in the planktonic fraction determined by qPCR of 16S rRNA genes (Total Bacteria), *amoA* genes (Comammox, AOB), 16S rRNA genes (Anammox), and *nxr* genes (NOB). Gene copy numbers per ng of total extracted DNA (n=3) are shown for samples taken at the start of the incubation (day 0) and after 86 days (from both beads and planktonic materials). Genetic material corresponding to the initial biomass (day 0) was obtained directly after immobilization through the same extraction method as 86 days biomass samples. Gene copy numbers were adjusted according to their average occurrence (Suppl. Table 3).

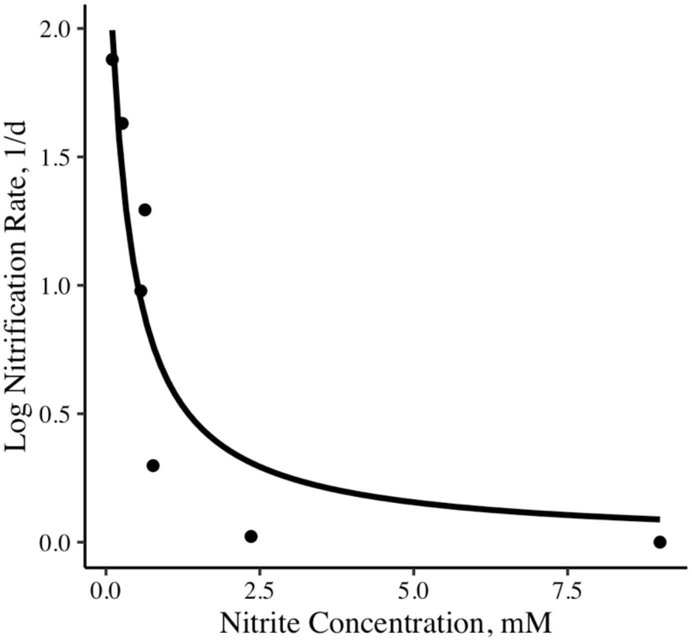


**Supplementary Figure 4.** Nitrite inhibition of *N. inopinata*. Each point corresponds to a batch culture of *N. inopinata* incubated at a given nitrite concentration in which the nitrification rate was inferred from the nitrate production rate. A typical inhibition curve was fitted to the data [16].
